# Aqp1aa and Aqp4a mediate collecting duct water permeability in a marine teleost

**DOI:** 10.64898/2026.05.16.725598

**Authors:** Erika Watanabe, Chihiro Ota, Genki Imaizumi, Yohei Sakamoto, Yutaka Suzuki, Akira Kato

## Abstract

Regulation of water permeability in the collecting duct is important for osmoregulatory acclimation in teleost fish. In hyperosmotic environments such as seawater (SW), the teleost kidney functions as a site of divalent ion excretion. The collecting ducts reabsorb Na^+^, Cl^−^, and water, thereby reducing urine volume and producing small amounts of isotonic urine with high concentrations of divalent ions. In hypoosmotic environments such as freshwater (FW) or low-salinity brackish water (BW), the kidney produces large volumes of hypotonic urine and serves as a site of water excretion; under these conditions, the collecting ducts reabsorb Na^+^ and Cl^−^ but not water. To identify aquaporins (Aqps) involved in regulating water permeability in the collecting ducts of teleosts, we analyzed renal Aqp expression in a euryhaline marine fish, the Japanese pufferfish (*Takifugu rubripes*), which possesses 16 Aqp genes in its genome, seven of which (Aqp1aa, 1ab, 3a, 4a, 7, 8bb, and 11a) are expressed in the kidney. Quantitative RT-PCR analysis showed that Aqp1aa and Aqp4a were highly expressed in collecting duct tissues, and that Aqp1aa expression was markedly reduced in fish acclimated to BW. Immunohistochemistry revealed apical localization of Aqp1aa and basolateral localization of Aqp4 in collecting duct cells, with apical Aqp1aa downregulated in BW. These results suggest that Aqp1aa and Aqp4 mediate water reabsorption in SW and that downregulation of Aqp1aa contributes to hypotonic urine production in BW.

**NEW & NOTEWORTHY:** Regulation of water permeability in the collecting duct is important for osmoregulation in teleost fish. Expression analyses of aquaporins (Aqps) in the marine pufferfish *Takifugu rubripes* showed that Aqp1aa and Aqp4a are highly expressed in the collecting duct and localized to the apical and basolateral membranes, respectively. Renal Aqp1aa expression was markedly reduced in fish acclimated to hypoosmotic brackish water. These results indicate that collecting duct water permeability is regulated by Aqp1aa expression.

## INTRODUCTION

The composition of extracellular fluids in teleost fishes is similar to that of humans and other tetrapods (1, 2). Teleosts maintain ion homeostasis of body fluids to acclimate to hyperosmotic (seawater, SW) or hypoosmotic (freshwater, FW; low-salinity brackish water, BW) environments. SW has an osmolarity approximately three times that of body fluids and contains high concentrations of Na^+^ and Cl^−^ as well as divalent ions. To compensate for water loss and ion gain, SW teleosts drink seawater, absorb ions and water through the intestine (3), and excrete excess ions via the gills and kidney (4, 5). The kidney produces small amounts of isotonic urine rich in divalent ions and serves as a site of divalent ion excretion (2, 6, 7). In contrast, FW has an osmotic pressure less than 1/100 that of body fluids. To counter water influx and ion loss, FW teleosts absorb salts from the environment via ionocytes in the gills (4, 5, 8). The kidney produces large volumes of hypotonic urine and serves as a site of water excretion (2, 7).

The collecting duct is the terminal segment of the renal tubule in the teleost kidney (7, 9, 10). In seawater (SW) teleosts, nephrons mainly consist of Bowman’s capsule, proximal tubule, and collecting duct, although some species lack glomeruli and are known as aglomerular fish. In these fish, primary urine is produced solely by tubular secretion (6). In SW teleosts, proximal tubules actively secrete fluid containing SO_4_ ^2−^ and Mg^2+^, and the collecting ducts subsequently reabsorb Na^+^, Cl^−^, and water, thereby reducing urine volume and producing small amounts of isotonic urine with high concentrations of divalent ions. In contrast, in freshwater (FW) or brackish water (BW) teleosts, nephrons generally consist of Bowman’s capsule, proximal tubule, distal tubule, and collecting duct, and primary urine is produced by glomerular filtration and tubular secretion (7, 9, 10). Under these conditions, glomeruli generate large volumes of filtrate, and the collecting ducts reabsorb Na^+^ and Cl^−^ but not water, resulting in the production of large volumes of hypotonic urine (11, 12). Thus, water permeability of the collecting duct is thought to differ markedly between SW and FW/BW teleosts.

Regulation of water permeability in the collecting duct has been well studied in mammals. Aquaporin 2 (Aqp2) is localized to the apical membrane of collecting duct cells, and its cell surface expression is regulated by antidiuretic hormone (ADH) via receptors expressed in these cells (13, 14). Upon ADH stimulation, Aqp2 translocates from intracellular vesicles to the apical membrane, whereas inhibition of ADH secretion leads to its internalization. In mammals, *aqp2* is located in tandem with *aqp5* and *aqp6* on the same chromosome (15). Comparative analyses of vertebrate aquaporins indicate that teleosts lack *aqp2*/*5*/*6*. Therefore, teleosts likely regulate collecting duct water permeability by using other Aqp(s), but the underlying molecular mechanisms remain unclear.

To investigate these mechanisms, we focused on an euryhaline marine teleost, the Japanese pufferfish (*Takifugu rubripes*). Previous studies have shown that its genome contains 16 *aqp* genes, seven of which are expressed in the kidney (16). In the present study, we identified two aquaporins, Aqp1aa and Aqp4a, that are highly expressed in the collecting ducts. In SW-acclimated fish, Aqp1aa and Aqp4a are localized to the apical and basolateral membranes, respectively, whereas Aqp1aa expression is markedly downregulated in fish acclimated to BW. These findings provide insight into the molecular mechanisms regulating water permeability in the collecting ducts of teleost fish.

## MATERIALS AND METHODS

### Animals

Marine pufferfish (Japanese pufferfish, torafugu, *Takifugu rubripes*) were purchased from a local dealer, reared in natural seawater (SW, ∼35 ppt) for more than 7 days, and then transferred to 150-L tanks containing 1-ppt brackish water (1-ppt BW, ∼3% diluted natural SW) for 8–9 days (BW torafugu) as described previously (17). The experimental animals were anesthetized by immersion in 0.1% ethyl m-aminobenzoate (MS-222, tricaine; Sigma-Aldrich, St Louis, MO, USA) neutralized with sodium bicarbonate. After humane killing by cervical transection, the tissues were dissected. Pufferfishes were housed and cared in accordance with a manual approved by the Institutional Animal Experiment Committee of the Institute of Science Tokyo.

Isolation of total RNA from the whole kidney and the collecting duct of pufferfish Immediately after a pair of kidneys were removed from pufferfish, one kidney was sliced ∼ 5 mm thick and immersed in RNAlater solution (Thermo Fisher Scientific, Waltham, MA, USA). This preparation was used to isolate total RNA from the whole kidney. The other kidney was immediately immersed in RNAlater solution, and the collecting ducts were dissected using a stereomicroscope and tweezers. Collecting ducts were distinguished from other tissues by shape and diameters (200 – 800 µm) (11, 18) and pooled to isolate total RNA from the collecting ducts. Total RNAs of the pufferfish whole kidneys and the collecting duct tissues were isolated using the acid guanidinium thiocyanate–phenol–chloroform extraction method with Isogen (Nippon Gene, Tokyo, Japan). The concentration and quality of RNA were measured based on UV absorbance at 260 and 280 nm and checked using denaturing agarose gel electrophoresis or Microchip Electrophoresis System for DNA/RNA Analysis MCE-202 MultiNA (Shimadzu, Kyoto, Japan) with an RNA reagent kit (Shimadzu).

### Quantitative PCR (qPCR) analysis

First-strand complementary DNAs were synthesized from 5 μg of total RNA using the SuperScript IV First-Strand Synthesis System (Thermo Fisher Scientific) with oligo(dT) primers, diluted eightfold with nuclease-free water, and used as a template for qPCR analysis. The relative expression of *aqp1aa, aqp1ab, aqp3a, aqp4a, aqp7, aqp8bb*, and *aqp11a* was analyzed by quantitative real-time PCR (qPCR) using the SYBR Green method, with *actb* as reference genes, and primers listed in Table 1. For qPCR analysis, TB Green Premix Ex Taq II (Takara Bio) and the Thermal Cycler Dice Real Time System III (Takara Bio) were used. The statistical significance set at *p* < 0.05 was evaluated by one-way analysis of variance (ANOVA) followed by the Tukey–Kramer multiple comparisons test GraphPad Prism software (Version 5, GraphPad, San Diego, CA). Origin software (Version 8, OriginLab, Northampton, MA) was used to display the results in box plots.

**Table 1.**
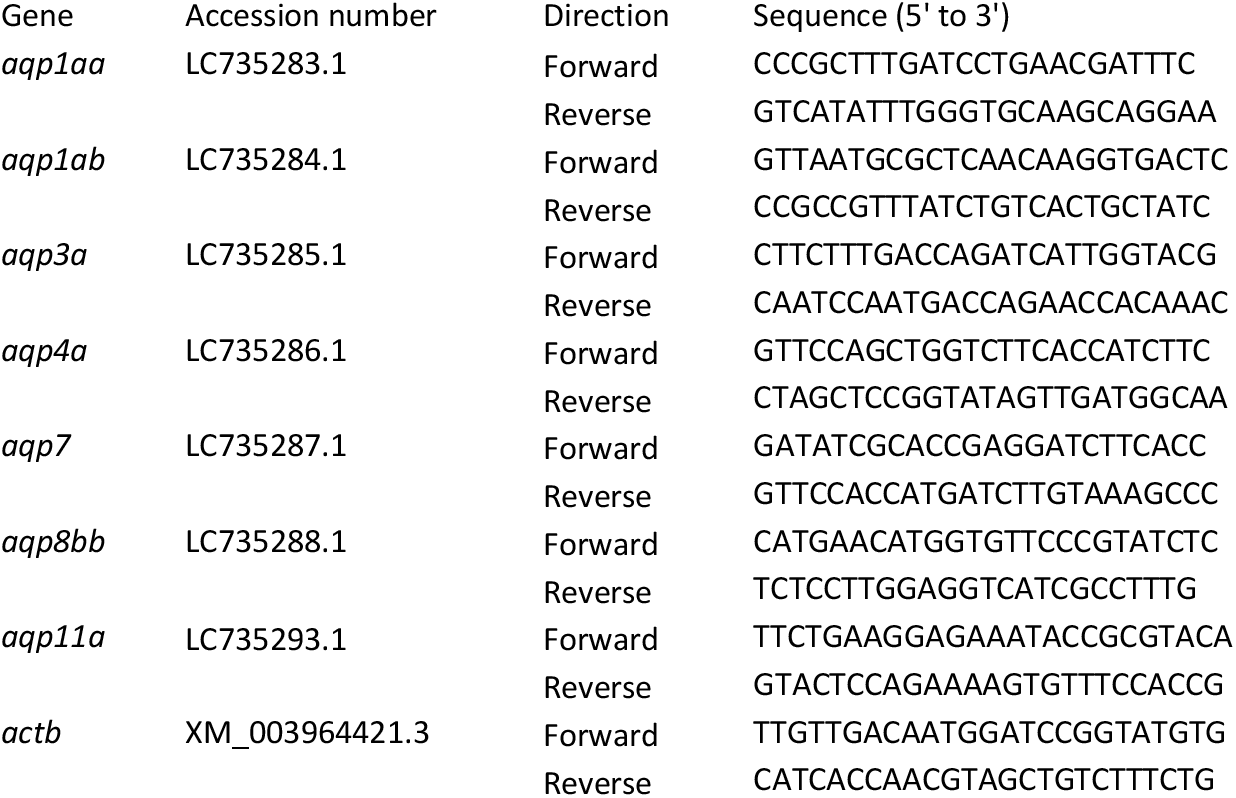
Primers used for quantitative PCR in this study.

### Antibody Preparation

Polyclonal antisera were made in rabbits that had been immunized with BSA-conjugated synthetic peptides corresponding to parts of Japanese pufferfish Aqp1aa (DDBJ/EMBL/GenBank accession number LC735283, amino acid residues 225–261, PKYDDFPERMKVLVSGPVGDYDVNGGNDNTTVEMTSK) or Aqp4a (LC735286, amino acid residues 236–282, ELKKRLETVFHKDSAGRYREVEAEDVAIKPGSVHTVSHLEKAEKKEC; amino acid residues 282–294, CFQDTAGEVLSSV).

Antibody specificity was established by staining CHO cells exogenously expressing pufferfish Aqp1aa and Aqp4a. Full-length cDNAs of Aqp1aa and Aqp4a were isolated from the kidney of Japanese pufferfish (*Takifugu rubripes*) by RT-PCR using previously prepared first-strand cDNA (19), the primers designed based on the cDNA sequence (DDBJ/EMBL/GenBank accession numbers LC735283 and LC735286) and a high-fidelity DNA polymerase (KOD-Plus-Neo DNA polymerases, Toyobo, Osaka, Japan). The primer sequences are: 5’-attcatcgatagacatgagagagttgaagagcaag-3’ and 5’-cgactggtaccgatatttttgatgtcatctcc-3’ for Aqp1aa; 5’-attcatcgatagacatgtgtggctgggcgtccctc-3’ and 5’-cgactggtaccgatattacgggaggacagcacttc-3’ for Aqp4a. The amplified DNA was subcloned into p3×FLAG-CMV-14 for the expression of C-terminal 3xFLAG-tagged Aqps in mammalian culture cells.

CHO cells were grown in Ham’s F-12 nutrient mixture (F-12) with 10% FBS, 100 U/mL penicillin, and 100 µg/mL streptomycin at 37 °C in a humidified 5% CO_2_ atmosphere, cultured on coverslips in 12-well plates, and transfected with plasmids p3×FLAG-CMV-14-Aqp1aa, p3×FLAG-CMV-14-Aqp4a using Lipofectamine LTX (Invitrogen), as described previously (11, 17). For immunofluorescence experiments, at 36 h after transfection, the transfected and untransfected cells were fixed and stained with anti-Aqp1aa or anti-Aqp4a rabbit sera (1:1,000 dilution) mixed with mouse M2 anti-FLAG monoclonal antibody (3.5 μg/ml, Sigma-Aldrich) for 1 h at room temperature, as described previously (11). Cells were then treated with Alexa Fluor 488-labeled F(ab’)2 fragment of donkey anti-rabbit IgG (1:2,000; Jackson ImmunoResearch Laboratories, West Grove, PA, USA), Cy3-labeled F(ab’)2 fragment of goat anti-mouse IgG (1:2,000; Jackson ImmunoResearch Laboratories), and Hoechst 33342 (100 ng/ml; Thermo Fisher Scientific) for 1 h at room temperature. Fluorescence images were obtained using a confocal laser scanning microscope (CLSM, LSM780; Carl Zeiss, Oberkochen, Germany) and processed using ZEN software (Carl Zeiss). The images of the test samples and negative controls were acquired under identical settings.

### Immunohistochemistry

Immunohistochemical analyses of frozen sections (6 μm) of the Japanese pufferfish kidney were performed, as described previously (11, 17). The sections were incubated with anti-Aqp1aa, anti-Aqp4a, or corresponding preimmune rabbit sera (1:1,000 dilution) together with anti-Na^+^/K^+^-ATPase (Nka) rat serum (1:1,000) or nonimmune rat serum (1:1,000 dilution) (11, 20) for 8 h at room temperature. Sections were then washed with phosphate buffered saline and stained with Alexa Fluor 488-labeled F(ab’)2 fragment of donkey anti-rabbit IgG (1:2,000), Cy3-labeled F(ab’)2 fragment of goat anti-rat IgG (1:2,000; Jackson ImmunoResearch Laboratories), and Hoechst 33342 (100 ng/ml). Fluorescence images were acquired with a fluorescence microscope (model BZ-X1000; Keyence, Osaka, Japan). The images of the test samples and negative controls were acquired under identical settings.

## RESULTS

### Isolation of collecting duct tissue and evaluation

Observation of the Japanese pufferfish kidney tissue sections showed that the diameter of the proximal tubules is ∼50 um and the diameter of the collecting ducts is approximately 200–800 μm (11, 18). Since a nephron of Japanese pufferfish consists mainly of Bowman’s capsule, proximal tubule, and collecting duct, the tubular tissue with a diameter of 200 um or more obtained from Japanese pufferfish kidney tissue was expected to be collecting ducts. A preliminary RNA-Seq analysis of the collecting duct tissues isolated from a Japanese pufferfish showed that the expression of marker genes, *nkcc2*, was increased approximately 10-fold in the collecting duct compared to the whole kidney (data not shown), suggesting that collecting duct tissues were correctly isolated in this method (data not shown) (17).

Next, collecting duct tissues were isolated from five each of Japanese pufferfish acclimated in SW and BW (10 fish in total) using the same method, and RNAs were extracted. The expression levels of *nkcc2, clcnk*, and *slc22a2* were quantified by qPCR (Fig. 1A). The expression levels of *nkcc2* and *clcnk*, markers of collecting ducts (11, 21), were approximately 6-to 13-fold more enriched in the collecting ducts than in the kidney, although most comparisons were not statistically significant. The expressions of *slc22a2*, a marker of proximal tubules, in the collecting ducts was ∼1/30 lower than those in the kidney. These results suggest that the obtained collecting duct tissue is useful for the analysis of the collecting duct-specific gene expression.

**Figure 1.**
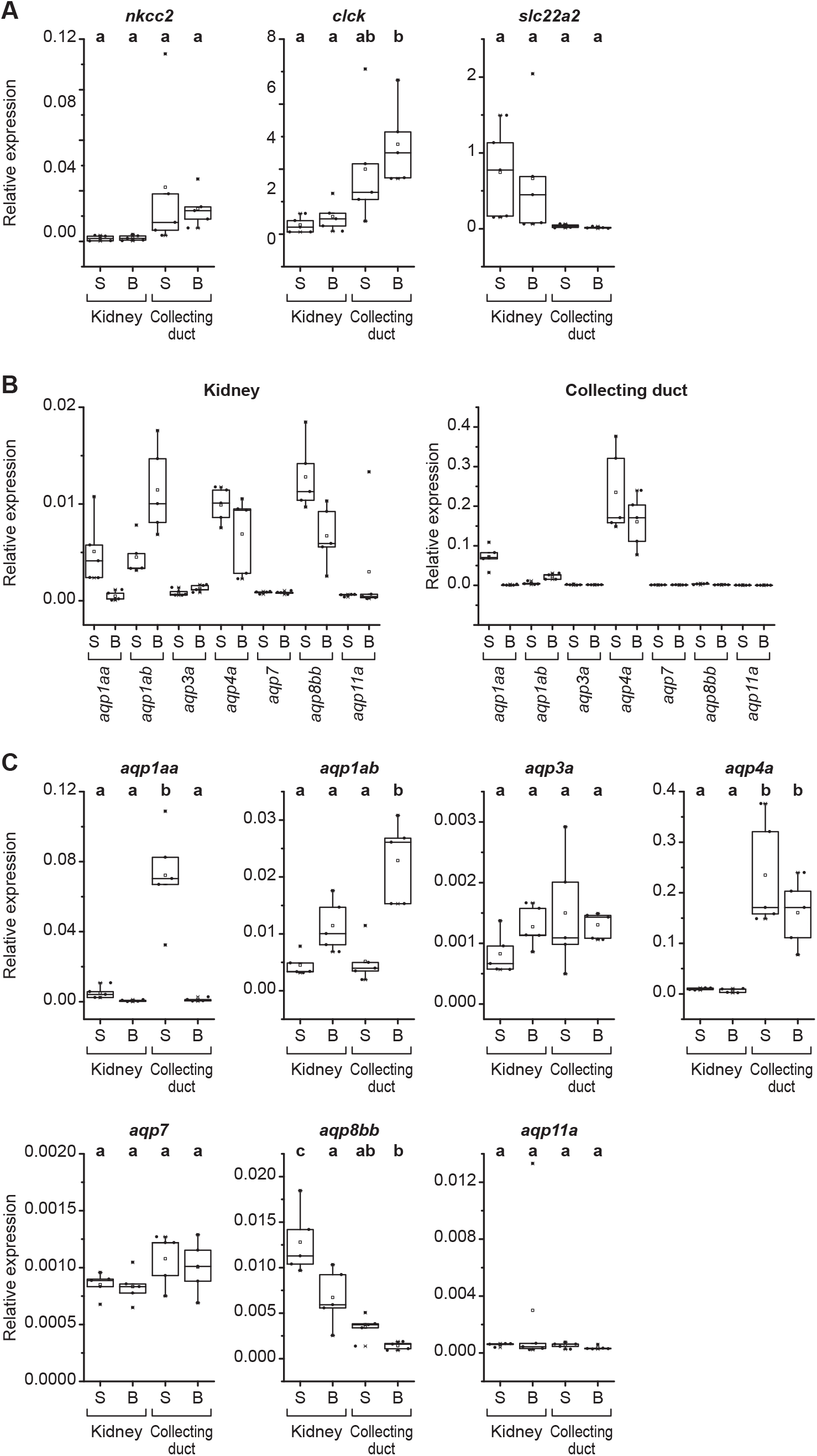
Quantitative PCR analysis of aquaporins in the whole kidney and collecting ducts of the SW- and 1-ppt BW acclimated Japanese pufferfish. The relative expression levels normalized to *actb* are presented as interquartile range from the 25–75 percentiles (box), range (whiskers), outliers (>1.5× interquartile range above upper quartile), mean (square in the box), and median (line in the box). (*A*) Quantitative PCR analysis of collecting duct markers (*nkcc2, clck*) and a proximal tubule marker (*slc22a2*). Different letters indicate significant differences (*p* < 0.05), whereas groups sharing the same letter are not significantly different (Tukey–Kramer test). (*B*) Quantitative PCR analysis of aquaporins. (*C*) Graphs summarizing the data in (*B*) by gene, with results of statistical analyses. Different letters indicate significant differences (*p* < 0.05), whereas groups sharing the same letter are not significantly different (Tukey–Kramer test).

### Expression of the *aqp* gene family in the kidney and collecting ducts of SW-acclimated Japanese pufferfish

In seawater (SW), the natural habitat of the Japanese pufferfish, the osmolarity of bladder urine was ∼390 mOsm/kg H2O, which is slightly lower but comparable to that of plasma (∼470 mOsm/kg H2O) (17). Previous studies have shown that the kidney of the Japanese pufferfish expresses seven aquaporin genes (*aqp*): *aqp1aa, aqp1ab, aqp3a, aqp4a, aqp7, aqp8bb*, and *aqp11a* (16). To identify which *aqp* genes are abundantly expressed in the collecting ducts, we compared their expression levels between the kidney and isolated collecting duct tissues using quantitative real-time PCR (Fig. 1B). In the kidney, *aqp1aa, aqp1ab, aqp4a*, and *aqp8bb* were highly expressed, whereas *aqp3a* and *aqp11a* were expressed at low levels. In the collecting ducts, *aqp1aa* and *aqp4a* were highly expressed. Notably, in SW-acclimated fish, the expression levels of *aqp1aa* and *aqp4a* in the collecting ducts were approximately 14- and 24-fold higher than in the whole kidney, respectively. These results suggest that *aqp1aa* and *aqp4a* are the major aquaporin genes expressed in the collecting ducts of the Japanese pufferfish.

### Changes in expression of the *aqp* gene family in the kidney and collecting ducts of BW-acclimated Japanese pufferfish

In 1-ppt brackish water (BW), a hypoosmotic environment, the osmolarity of bladder urine was ∼40 mOsm/kg H2O, which is approximately one-eighth that of plasma (∼330 mOsm/kg H2O) (17). We examined changes in the expression of *aqp* genes in the kidney and collecting ducts of Japanese pufferfish acclimated to 1-ppt BW. Notably, expression of *aqp1aa* in both the collecting ducts and the whole kidney decreased to approximately one-seventieth and one-tenth of the levels observed in SW-acclimated fish, respectively, whereas *aqp4a* expression did not change significantly (Fig. 1C). In contrast, expression of *aqp1ab* in the kidney increased during 1-ppt BW acclimation (Fig. 1C).

### Evaluation of anti-Aqp1aa and anti-Aqp4a antibodies

Peptides corresponding to partial sequences of Japanese pufferfish Aqp1aa and Aqp4a were synthesized, and polyclonal antibodies were raised in rabbits against these peptides. To evaluate the specificity of the anti-Aqp1aa and anti-Aqp4a antibodies, C-terminally 3×FLAG-tagged proteins (Aqp1aa-3×FLAG and Aqp4a-3×FLAG) were expressed in CHO cells and analyzed by immunofluorescence. Anti-FLAG antibody produced strong signals, confirming the expression of Aqp1aa-3×FLAG and Aqp4a-3×FLAG in CHO cells (Fig. 2). Anti-Aqp1aa and anti-Aqp4a antibodies showed staining patterns similar to those observed with the anti-FLAG antibody (Fig. 2). No staining was detected in untransfected CHO cells stained with anti-Aqp1aa, anti-Aqp4a, or anti-FLAG antibodies. These results indicate that the anti-Aqp1aa and anti-Aqp4a antibodies specifically recognize Aqp1aa and Aqp4a, respectively.

**Figure 2.**
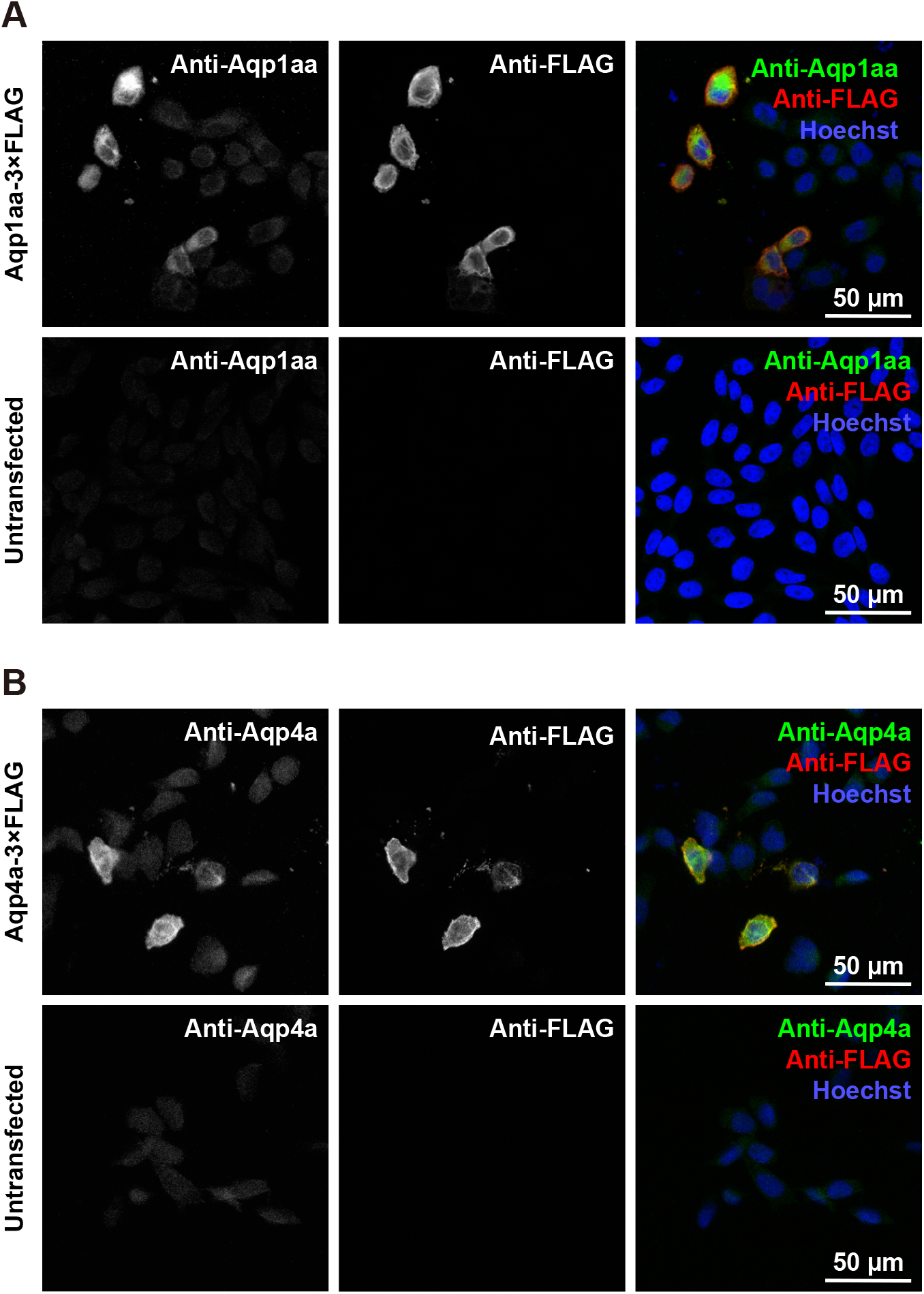
Validation of the specificity of anti-Aqp1aa and anti-Aqp4a antibodies. (*A*) Immunostaining of CHO cells transfected with an Aqp1aa-3×FLAG expression vector or untransfected cells, using rabbit anti-Aqp1aa (green) and anti-FLAG (red) antibodies. (*B*) Immunostaining of CHO cells transfected with an Aqp4a-3×FLAG expression vector or untransfected cells, using rabbit anti-Aqp4a (green) and anti-FLAG (red) antibodies. Nuclei were stained with Hoechst 33342 (blue). Scale bars, 50 μm.

### Localization of Aqp1aa and Aqp4a in the kidney of Japanese pufferfish

The localization of Aqp1aa and Aqp4a in the kidney of Japanese pufferfish was analyzed by immunofluorescence (Fig. 3, left pannels). In SW-acclimated fish, anti-Aqp1aa and anti-Aqp4a antibodies labeled the apical and basolateral regions of collecting duct cells, respectively. Anti-Na^+^ /K^+^ -ATPase (Nka) antibody was used as a basolateral membrane marker (11). No signals were detected with preimmune or nonimmune sera. These results indicate that, in the collecting ducts of SW-acclimated fish, Aqp1aa is localized to the apical membrane, whereas Aqp4a is localized to the basolateral membrane.

**Figure 3.**
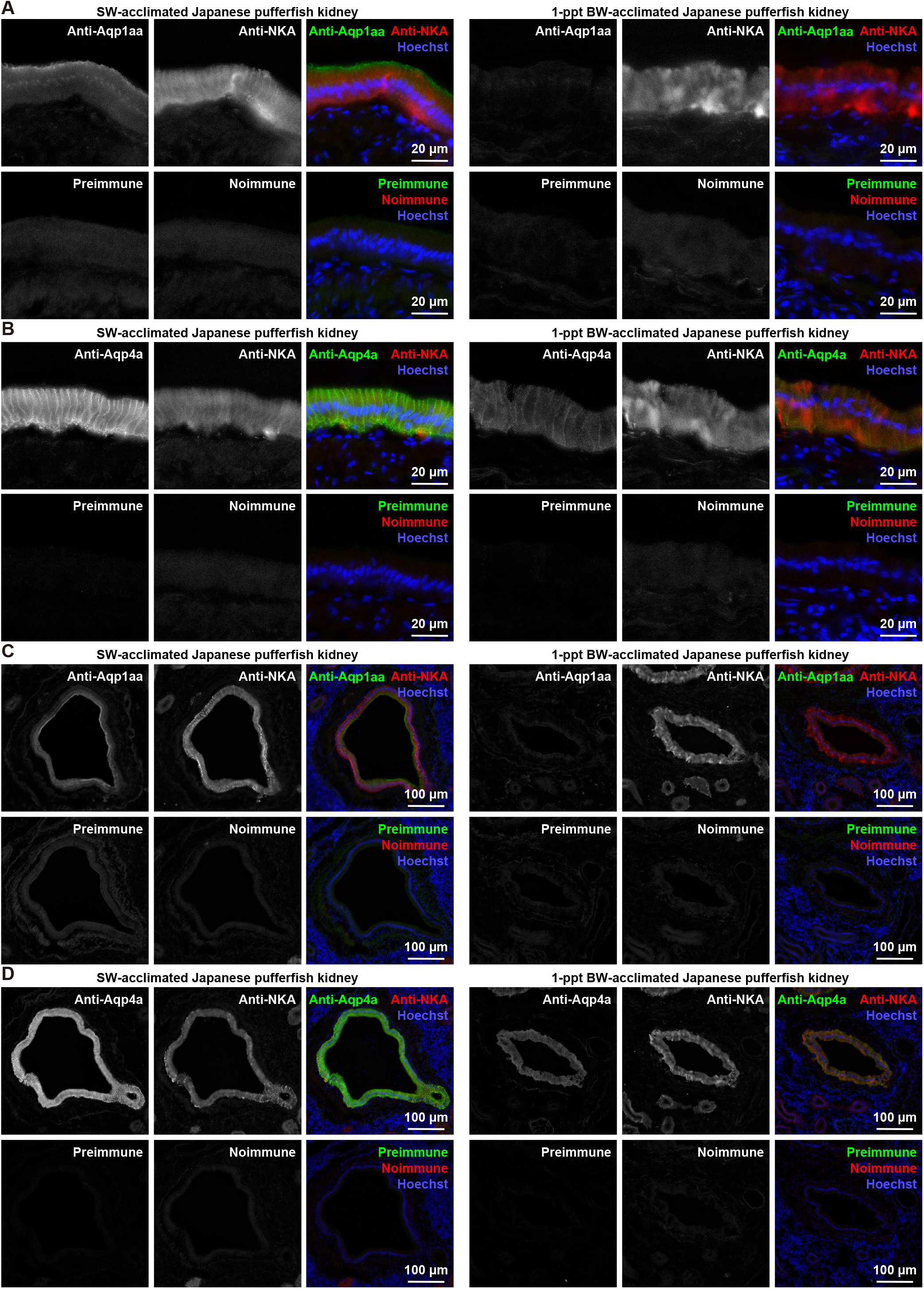
Localization of Aqp1aa and Aqp4a in the kidneys of SW- and 1-ppt BW–acclimated Japanese pufferfish. (*A*) High-magnification images of collecting ducts stained with anti-Aqp1aa (green) and Nka (red) antibodies. (*B*) High-magnification images of collecting ducts stained with anti-Aqp4a (green) and Nka (red) antibodies. (*C*) Low-magnification images of collecting ducts stained with anti-Aqp1aa (green) and Nka (red) antibodies. (*D*) Low-magnification images of collecting ducts stained with anti-Aqp4a (green) and Nka (red) antibodies. For negative controls, tissue sections were stained with preimmune rabbit serum and nonimmune rat serum. Nuclei were stained with Hoechst 33342 (blue).

The same analysis was performed in the kidneys of fish acclimated to 1-ppt BW (Fig. 3, right pannels). Anti-Aqp4a antibody labeled the basolateral region of collecting duct cells similarly to that observed in SW-acclimated fish. In contrast, anti-Aqp1aa antibody did not produce detectable signals at the apical membrane in BW-acclimated fish.

## DISCUSSION

SW teleosts produce isotonic urine rich in divalent ions, whereas FW teleosts produce hypotonic urine; these processes are fundamental to osmoregulatory acclimation (2, 7, 9, 10). The production of urine with different osmotic properties requires precise regulation of water permeability along the renal tubules. High water permeability facilitates water reabsorption, thereby reducing urine volume and enabling the production of isotonic urine, whereas low water permeability is essential for the production of hypotonic urine. In the present study, to elucidate the mechanisms regulating water permeability in the collecting duct, we examined the expression and localization of Aqps in the collecting ducts of a euryhaline marine teleost, the Japanese pufferfish, acclimated to SW or hypoosmotic 1-ppt BW. In SW-acclimated fish, *aqp1aa* and *aqp4a* were highly expressed in the collecting ducts, and immunofluorescence revealed that Aqp1aa and Aqp4a are localized to the apical and basolateral membranes, respectively. These findings suggest that Aqp1aa and Aqp4a contribute to maintaining high water permeability in the collecting duct, thereby facilitating water reabsorption and the production of isotonic urine in SW. In contrast, in fish acclimated to 1-ppt BW, *aqp1aa* expression was markedly reduced at the mRNA level, and immunofluorescence showed a loss of Aqp1aa signals at the apical membrane of collecting duct cells. The expression level and localization of Aqp4a remained largely unchanged. These results suggest that downregulation of Aqp1aa at both the transcript and protein levels reduces water permeability in the collecting duct under hypoosmotic conditions, thereby contributing to the production of hypotonic urine. These findings indicate that Aqp1aa expressed in the collecting duct is one of the key molecules involved in osmoregulation in teleost fish. Furthermore, these findings suggest that, in the Japanese pufferfish, a teleost lacking Aqp2, which mediates water permeability in the mammalian collecting duct, Aqp1aa may serve a function analogous to that of mammalian Aqp2.

Comprehensive analyses of *aqp* genes in more than 400 fish genomes by Ferré et al. revealed that teleosts possess a tandemly arranged *aqp1aa–aqp1ab2–aqp1ab1* gene cluster, which is specific to teleosts and absent in basal actinopterygians (22). In the Japanese pufferfish, one copy of the *aqp1ab* gene has been secondarily lost, resulting in a tandem arrangement of *aqp1aa* and a single *aqp1ab* gene. Functional studies using *Xenopus* oocyte expression systems have demonstrated that teleost Aqp1 paralogs exhibit water-selective permeability and lack permeability to solutes such as glycerol and urea (16, 23-25). In the teleost intestine, salinity-dependent changes in *aqp1* gene expression and apical localization of Aqp1 in epithelial cells have been reported (26, 27), indicating that Aqp1 mediates water transport across the apical membrane. In these cells, salt absorption via apical Nkcc2 and basolateral Na^+^ /K^+^ -ATPase drives water absorption (3, 28, 29). In the kidney, *aqp1aa* expression has been shown to be elevated under hyperosmotic conditions (SW) and reduced under hypoosmotic conditions in species such as sea bass (*Dicentrarchus labrax*) (30, 31) and river puffer (*Takifugu obscuru*s) (32). Consistent with these observations, the present study demonstrates that renal *aqp1aa* expression in the Japanese pufferfish is high in SW-acclimated fish and markedly reduced in fish acclimated to 1-ppt BW. These findings suggest that transcriptional regulation of *aqp1aa*, upregulated under hyperosmotic conditions and downregulated under hypoosmotic conditions, is conserved across teleost species. In contrast, in the European eel (*Anguilla anguilla*), expression of Aqp1 and its duplicate (Aqp1 dup) in the kidney is lower under SW conditions than under freshwater conditions, and immunohistochemical analyses have localized Aqp1 to the apical membrane of proximal tubules and to endothelial cells (33). This pattern differs from the present findings in the Japanese pufferfish, suggesting that the renal function of Aqp1aa may vary among species. Interestingly, expression of *aqp1ab* in the Japanese pufferfish was upregulated under 1-ppt BW acclimation, a pattern that appears similar to that observed for Aqp1 and Aqp1 dup in the European eel (33). Further detailed analyses of the localization of Aqp1ab in the pufferfish kidney will be necessary to clarify the functional relationships and potential divergence among these paralogs.

Teleosts possess two *aqp4* ohnologs, *aqp4a* and *aqp4b*, generated by the teleost-specific whole-genome duplication (15); however, in the Japanese pufferfish, one copy has been secondarily lost, and only *aqp4a* is retained (15, 16). Aqp4a has been confirmed to function as a water channel in teleosts (16, 23). In mammals, Aqp4 is localized to the basolateral membrane of collecting duct cells (13, 34). In contrast, the localization of Aqp4 in the kidney of cartilaginous fish has been examined in detail, and its role in urea reabsorption has been suggested (35, 36); however, no studies have investigated the localization of Aqp4 in the kidney of teleost fish. The present study demonstrates that Aqp4a is localized to the basolateral membrane of collecting duct cells in the Japanese pufferfish. This finding suggests that the function of Aqp4 in the collecting duct is conserved between mammals and teleosts. Furthermore, neither the expression level nor the localization of Aqp4a differed between SW and 1-ppt BW conditions. Given that the basolateral membrane is in direct contact with the interstitial fluid and provides the fundamental transport machinery necessary for cellular homeostasis, Aqp4a may contribute not only to water reabsorption under SW conditions but also to basal water transport in both SW and BW environments.

## DATA AVAILABILITY

Data will be made available upon reasonable request.

## ACKNOWLEDGMENTS

We thank Dr. Kohei Hosono, Keiichiro Fujimori, and Tomohiro Nakao for contribution at the early stage of this work, Dr. Ayumi Nagashima for discussion, Yoko Yamamoto, Nana Shinohara, the Biomaterials Analysis Division, and the Open Research Facilities for Life Science and Technology at the Institute of Science, Tokyo, for their technical assistance.

## AUTHOR CONTRIBUTIONS

E.W., C.O., G.I., Y.Sa., and A.K. conceived and designed research; E.W., C.O., G.I., Y.Su. and A.K. analyzed data; E.W., C.O., G.I., Y.Sa., Y.Su. and A.K. performed experiments; E.W., C.O., G.I., and A.K.; interpreted results of experiments; E.W., C.O., G.I., and A.K. prepared figures; E.W., C.O., G.I., and A.K. drafted manuscript; C.O. and A.K. edited and revised manuscript; E.W., C.O., G.I., Y.Sa., Y.Su. and A.K. analyzed data; E.W., C.O., G.I., Y.Sa., Y.Su. and A.K. approved final version.

